# From womb to crib: How fetal activity patterns *in utero* reveal postnatal sleep behavior

**DOI:** 10.1101/2024.12.04.626754

**Authors:** Andjela Markovic, Christophe Mühlematter, Christine Blume, Petra Zimmermann, Salome Kurth

**Affiliations:** Department of Psychology, University of Fribourg, Fribourg, Switzerland; Department of Pulmonology, University Hospital Zurich, Zurich, Switzerland; Department of Social Neuroscience and Social Psychology, University of Bern, Bern, Switzerland; Centre for Chronobiology, Psychiatric Hospital of the University of Basel, Basel, Switzerland; Research Cluster Molecular and Cognitive Neurosciences, University of Basel, Basel, Switzerland; Department of Biomedicine, University of Basel, Basel, Switzerland; Department of Pediatrics, Hôpital Fribourgeois, Fribourg, Switzerland

## Abstract

**Importance:** The human circadian system, a critical biological mechanism governing the sleep-wake cycle, begins to oscillate before birth. Distinct fetal behavioral states have been described using short-duration ultrasound recordings, but the alignment of fetal activity with day and night cycles (i.e., diurnal rhythms) remains poorly understood due to the lack of long-term monitoring. Accordingly, a substantial knowledge gap exists concerning the evolution of fetal diurnal rhythms into infant sleep-wake cycles.

**Objective:** To investigate the development of fetal diurnal rhythms and their evolution into postnatal infant sleep, as well as the role of maternal factors and melatonin in shaping early circadian entrainment.

**Methods:** In this cohort study, conducted in 2022 and 2023, we collected over 20 000 hours of wearable acceleration and temperature data from 32 fetuses and their mothers during continuous 5-day monitoring in the third trimester of pregnancy. The cohort was subsequently followed longitudinally from the prenatal period through 6 months postpartum, with infant sleep outcomes assessed at three key time points (first postnatal weeks, 3 months, and 6 months) using sleep diaries and questionnaires. The first postnatal assessment additionally included breast milk and infant stool sample collection.

**Results:** Multi-level modelling revealed early diurnal rhythms in fetuses that align with maternal activity and circadian rhythms. Random Forest analyses identified fetal day/night sleep duration ratio – a simplified proxy for fetal circadian entrainment – as the strongest predictor of postnatal infant sleep, with fetuses preferring nighttime sleep maintaining this preference postnatally. Maternal sleep regularity (i.e., consistent sleep patterns characterized by low day-to-day variability in sleep timing) during pregnancy also predicted infant sleep, associating with a stronger preference for nighttime sleep in infants.

**Conclusions and relevance:** These findings highlight the potential influence of intrauterine and maternal factors on the evolution of circadian entrainment. Our study contributes to a broader understanding of the earliest emergence of circadian rhythms, supporting future research on their long-term health impacts. Maternal sleep regularity stands out as the earliest modifiable target for interventions, offering an actionable pathway to promote infant circadian entrainment, with potential long-term benefits for family dynamics and well-being.

## Introduction

The human body’s circadian clock, a system governing most biological processes including the sleep-wake cycle, starts oscillating in utero ^1^. However, little is known about how these oscillations translate to diurnal patterns of sleep rhythms before birth and how they affect postnatal infant sleep behavior. Healthy sleep in early postnatal life likely plays an active role in core maturational processes such as cognitive development ^2^. On the other hand, poor sleep during this sensitive period of early childhood poses a risk for developing deficits in inhibitory control ^3^, emotional problems ^4^ and psychopathology later in life ^5^. Given the importance of sleep in early life and a lack of normative longitudinal data spanning from prenatal stages through infancy, the earliest development of sleep regulation and its determinants represent critical gaps in our knowledge. Furthermore, various studies underscore the profound impact of maternal health and lifestyle during pregnancy on infant health outcomes (e.g., ^6,7^). This reveals pregnancy as the earliest critical phase, both of vulnerability and opportunity for targeted interventions, thereby allowing the application of precision medicine approaches in clinical settings with the ultimate goal to improve infant care and long-term developmental outcomes.

Ultrasound allows to distinguish at least three fetal behavioral states during the last weeks of pregnancy: 1) quiet state with reduced movement and low heart rate variability, 2) active state with frequent bursts of movement and high heart rate variability, and 3) active state with continuous movement and regular fast heart rates ^8,9^. More recently, fetal electrocardiography (fECG) established a relation between these states and the maternal sleeping position in late pregnancy ^10^. Stone et al. ^10^ observed higher fetal activity if the mother was in a lateral position, whereas fetal quiet sleep was associated with a supine position. Furthermore, fetuses were more active in the first hours of maternal sleep as compared to the rest of the night ^10^, suggesting rhythmicity to be established by 36 weeks of gestation. Importantly, associations between fetal and postnatal behavioral state patterns have repeatedly been reported ^11^. For example, similar sleep stage profiles (i.e., the duration ratio of different sleep states) have been observed between the fetal and the early postnatal periods ^12^. Furthermore, intra-fetal alignment between physiological (heart rate) and behavioral (movement) measures reflecting regular patterns in biological processes can predict early postnatal outcomes like emotional and attentional regulatory capacity ^13^. Taken together, these observations imply a trait-like consistency in activity patterns around birth. Nevertheless, the lack of long-term observational data on fetal behavioral states and a clear connection between these patterns and postnatal outcomes hinder the translation to clinical recommendations.

As none of the aforementioned technologies can be applied continuously over a longer period of time, so far it has been challenging to investigate fetal diurnal rhythms. Recent advances in wearable technology have provided new avenues for continuous monitoring of physiological parameters via non-invasive sensing technology ^14^. Actigraphy, one of the first wearable modalities to be commercially implemented, records movement patterns by means of acceleration sensors ^15^. For example, wrist-worn accelerometers are suitable to inform users about their physical activity (i.e., step counts), and algorithms are used to deduce information about sleep rhythms. In sleep research, actigraphy is an established method to acquire data related to objective sleep behavior ^16^ across longer periods of time in a natural environment (for an example, see ^17^). We successfully applied this technology in infancy, leading to a comprehensive quantitative database of infant sleep rhythms ^18^. Such studies demonstrate the potential of actigraphy for scalable, population-based, at-home use, which renders it ideal both for clinical practice and long-term developmental studies. Building on this foundation, we have adapted actigraphy to overcome the limitations of traditional methods such as ultrasound and fetal ECG, which are restricted to short-term monitoring. Our approach enables continuous monitoring of fetal activity over extended periods, providing unprecedented insights into fetal sleep rhythms.

The development of the fetal circadian system, first observed in rodents by Reppert and Schwartz ^19^, is strongly influenced by the maternal circadian system. Animal studies, where the maternal circadian rhythm is disrupted by manipulating the environmental conditions such as light, show that such disruptions alter fetal circadian development (reviewed in ^20,21^). A key pathway for this communication is maternal melatonin ^22,23^, a neuroendocrine marker of the internal biological night whose secretion starts 2-3 hours before habitual bedtime ^24,25^. Maternal melatonin is transferred to the fetus, as evidenced by its recovery from fetal brains in rodents ^26^, where it influences the fetal circadian clock ^27^. By aligning fetal circadian rhythms with the external environment, maternal melatonin prepares the fetus for postnatal light-dark cycle ^28^. Importantly, a neuroprotective role of melatonin has been demonstrated, suggesting further relevance beyond sleep regulation ^29^. Research in rodents has linked circadian disruption in the fetus to later-life health issues including cardiovascular disease ^30^ and mood disorders ^31^. Even postnatally, infants remain exposed to maternal time cues via breast milk ^32^, while developing their own rhythmicity. Understanding the complex pathways by which maternal circadian rhythms influence fetal development can thus inform future research and clinical approaches in pediatrics, neurology, and chronobiology, with implications for improving long-term health outcomes.

In this study, we employed wearable acceleration sensors to continuously track fetal movement patterns from the abdomen of pregnant women over multiple days. Throughout this timeframe, we evaluated maternal activity and skin temperature to quantify their influence on fetal activity patterns. Subsequently, we conducted follow-up assessments with the families at several postnatal intervals - during the early postnatal period (i.e., 2 to 54 days), and at 3 and 6 months - to investigate the developmental dynamics of infant sleep. Additionally, we undertook continuous collection of infant stool and maternal breast milk samples over several days during the early postnatal assessment to examine diurnal fluctuations in melatonin levels as a possible mechanism regulating newborn sleep. Beyond observing diurnal rhythms in fetal activity patterns, we hypothesized that maternal rhythms and melatonin levels in breast milk and infant stool might play a role in establishing robust sleep-wake cycles in infants.

## Methods

### Study design

Thirty-two healthy women in the last trimester of pregnancy were enrolled in the study. Participants were recruited through maternity wards, midwives, pediatricians, obstetricians, social media, personal contacts, and flyers distributed in public places in the German-speaking parts of Switzerland. Each woman was screened for eligibility during a telephone interview. Exclusion criteria were fetal malformations and chromosomal anomalies, intrauterine infections, maternal drug or alcohol use during pregnancy, preeclampsia, maternal travel across more than two time zones within the 30 days prior to the assessment and maternal sleep disorders. Written consent was obtained, and participants were informed of their right to discontinue participation at any time. The study was conducted in accordance with the Declaration of Helsinki and approved by the responsible ethics committee (*cantonal ethics committee*, BASEC 2019-02250). Families received small gifts as a compensation for their participation.

#### Pregnancy assessment

Between 28 and 40 weeks of gestation, participants continuously wore 7 sensor devices for at least 5 consecutive days to monitor physiological parameters (Figure 1). Four acceleration sensors (SENSmotion, SENS Innovation ApS, Copenhagen, Denmark) were placed on the abdomen to record abdominal wall deflections with a sampling rate of 25 Hz. Sensors were attached using dedicated patches to maintain their position and ensure comfort. One control acceleration sensor was placed on the chest to detect maternal movement. One participant wore only two acceleration sensors on the abdomen and a control acceleration sensor on the chest with a sampling rate of 12.5 Hz. One temperature sensor (iButton, Maxim Integrated, San Jose, USA) was placed under the right clavicle to record the participants’ temperature once per minute. Finally, one acceleration sensor worn on the left wrist was used to monitor maternal sleep as well as temperature with a sampling rate of 30 Hz (GENEActiv, Activinsights Ltd, Kimbolton, UK). Due to technical issues with the iButton temperature sensor, only temperature data recorded by the GENEActive sensor were analyzed.

**Figure 1:**
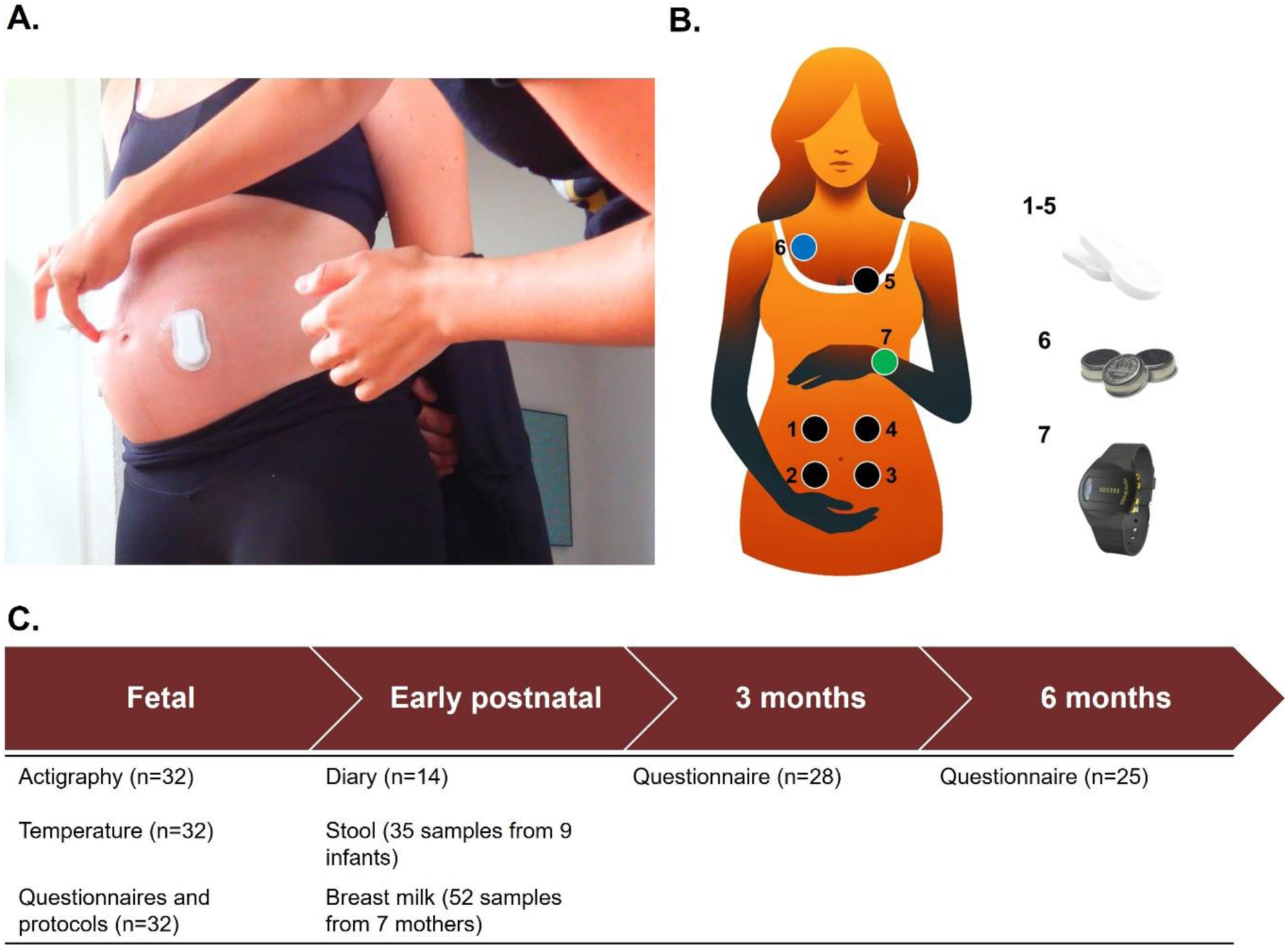
A. Home visits were conducted to provide participants with materials, attach sensors, and explain the study procedures. B. The placement of sensors is illustrated by circles, including four abdominal acceleration sensors (1 to 4, SENSmotion, SENS Innovation ApS, Copenhagen, Denmark), one chest acceleration sensor (5, SENSmotion, SENS Innovation ApS, Copenhagen, Denmark), one temperature sensor (6, iButton, Maxim Integrated, San Jose, USA) and one wrist acceleration and temperature sensor (7, GENEActiv, Activinsights Ltd, Kimbolton, UK). C. The timeline of the study shows the data types collected at different time points and the corresponding sample sizes in parentheses. Illustration in B created using DALL·E by OpenAI.

During these 5 days, participants filled out an activity protocol describing their activity at a 15-minute resolution based on four intensity categories (i.e., rest, sitting or standing activity, walking, sports) as well as their daily wake-up and bedtime. Additionally, during a 2-hour rest on each of the assessment days, they completed a minute-by-minute protocol marking the time intervals they perceived fetal movement. These two hours of daily rest were scheduled such that each session occurred at a different time of day across the 5 days, ensuring comprehensive coverage of various times throughout the day by the end of the assessment period. Finally, prior to giving birth, they completed an online questionnaire providing information on family background, health, and demographics.

#### Early postnatal assessment

Participants were re-contacted in the first weeks after birth to enroll for participation in the first postnatal assessment. Fourteen families agreed to participate in this part of the study and filled out a modified version of the Baby’s Day Diary ^33^. Over 5 consecutive days, they described their infant’s sleep, crying, and feeding behavior at a 15-minute resolution across 24-hour periods. Of these, 9 families collected samples (i.e., approximately 5 spatulas provided in 15 mL Sarstedt stool tubes) from each of their infant’s stools across a period of at least 24 hours documented in the diary. Additionally, mothers collected approximately 5 mL of breast milk into 15 mL Sarstedt tubes following each of the infant’s feedings. Two of the 9 mothers did not breastfeed their child, resulting in 7 datasets for breast milk. The samples were temporarily stored in the families’ freezer at -18°C and then transported to the laboratory in cooling boxes filled with crushed ice, typically within one month of collection. Upon arrival, samples were immediately stored at -80°C.

#### Follow-up survey

When the infants reached ages 3 and 6 months, participants received a link to an online questionnaire. This included questions about the mode of birth, the infant’s current weight and height, as well as their weight and height at birth, in addition to the Brief Infant Sleep Questionnaire ^34^. Data from participants who completed the questionnaire more than 1.5 months after their infant reached 3 or 6 months of age were excluded from the analysis to ensure accurate reflection of the specific developmental stages (see Figure 1 for analyzed sample size).

### Processing

Abdominal acceleration data were processed based on a protocol developed by Nishihara et al. ^35^ to detect fetal movement. Tri-axial (i.e., x, y and z axes) acceleration data were merged into one dimension as the square root of the sum of the squares of the three axes. The resulting signal was high-pass filtered at 0.5 Hz and low-pass filtered at 2.5 Hz. Integral amplitude was calculated across 50-ms intervals, except for the two sensors with a sampling rate of 12.5 Hz in one participant where the integral amplitude was calculated for 100-ms intervals. The remaining processing steps were kept consistent. Next, data from the control sensor was used to filter out artifacts caused by maternal movement from the abdominal sensor data. Specifically, peaks in chest acceleration signals were detected and removed from abdominal signals. Only data from maternal rest and sleep periods were included in further analyses amounting to an average of 50 hours of data per participant (i.e., 5 nights with approximately 8 hours of sleep and 5 days with 2 hours of daily rest). On these data, we applied a threshold set at the 75th percentile to the individual average of each 1-minute interval. If the average value exceeded the threshold, the interval was labelled as fetal activity; otherwise, it was classified as fetal inactivity.

Subsequently, we validated the extracted labels (i.e., fetal activity as 1 and fetal inactivity as 0) against maternal perception as reported in the diaries (i.e., reported fetal movement as 1 and missing reports as 0) using a moving Jaccard index ^36^, a measure of similarity defined as the ratio of intersection over union between two sample sets (i.e., labelled abdominal signals and maternal perception). We compared the labels within a moving window of 3 minutes to account for potential shifts in the noted clock time in the diaries. Finally, the calculated indices were averaged across all maternal rest periods to yield one value for each abdominal sensor. For each participant, the sensor with the highest average Jaccard index was selected for further analyses.

Maternal wrist acceleration data from the three axes were merged by using the square root of the sum of their squares, then bandpass-filtered between 3 and 11 Hz, and converted into activity counts per 15 seconds, following standard procedures ^37^. Subsequently, activity counts were summed for each minute and categorized as periods of sleep or wake using the algorithm developed by Sadeh et al. ^38^. From these categories, the sleep regularity index (SRI) was calculated based on the most frequently scored state across days for every minute ^39^. This measure represents the probability of being in the same state at a given time, providing an indicator of rhythm stability. Temperature data were cleaned using an automated procedure, which involved excluding all data points that were more than two standard deviations away from the mean, as previously applied to the same type of signals^40^. Missing values were filled using linear interpolation. Next, we averaged the temperature values for each 1-hour interval and collapsed to a 24-hour temperature curve. To estimate the phase of the circadian temperature rhythm, we used nonlinear least squares to fit a cosine function with a 24-hour periodicity to the resulting temperature curves, in line with previous work ^41^. The extracted temperature phase reflects the number of hours after midnight at which the temperature peaks. At this stage, periods when the devices were not worn were excluded.

### Melatonin analysis

Melatonin concentration in breast milk and stool was determined with a radioimmunoassay, RK-MEL2, manufactured by NovoLytiX GmbH (Witterswil, Switzerland) according to the instructions for use (IFU, version 2022-03-07). For each milk sample, 1 mL of breast milk was mixed with 1 mL of chloroform. The mixture was vortexed for 10 seconds at 3000 rpm, left for 10 minutes and vortexed again for 10 seconds at 3000 rpm. The mixture was then centrifuged for 10 minutes at 7000 x g in an Eppendorf Minifuge. 0.7 mL of the chloroform phase was then transferred into a borosilicate glass tube, and the chloroform was evaporated until dryness using a SpeeVac concentrator. The residues containing melatonin were resuspended with 1 mL of RK-MEL2 Incubation Buffer. Final concentrations were multiplied with a factor of 1.42 to correct for the loss resulting from using 0.7 mL of the 1 mL chloroform phase. For each stool sample, 50 mg of stool was homogenized in 1 mL of RK-MEL2 Incubation Buffer. The mixture was vigorously vortexed for 10 seconds and left for 10 minutes at room temperature. The latter step was repeated 3 times, and the homogenate was centrifuged for 10 minutes at 7000 x g in an Eppendorf Minifuge. The supernatant was transferred into a fresh tube and centrifuged again for 10 minutes at 7000 x g. The clear supernatant was then directly used for the immunoassay. The final concentrations were multiplied with factor of 0.77 to correct for the overestimated average recovery of 129% due to stool matrix effects (confirmed by spiking recovery experiments).

### Statistical analysis

#### Pregnancy time-series modelling

In the first part of our statistical analysis, we examined the association between maternal and fetal rhythms using the time-series of all signals collected during the pregnancy assessment and multi-level modelling. Z-scores were calculated to standardize all physiological signals and used as input to the multi-level models. We compared the performance of four models with regards to their Akaike Information Criterion (AIC) and root mean squared error (RMSE) with abdominal sensor measurements as the outcome variable. The four models included: 1) only a random intercept; 2) gestational age (in days), maternal vigilance state (i.e., wake/sleep with sleep serving as the reference state), maternal wrist skin temperature, maternal wrist activity, twin pregnancy (yes/no), in-vitro fertilization (yes/no), gestational diabetes (yes/no) and nulliparity (yes/no) as fixed factors; 3) random intercept and slope in addition to the fixed factors from 2); and, 4) interactions between gestational age, maternal vigilance state, maternal wrist skin temperature and maternal wrist activity in addition to all terms from 3). These fixed factors were selected based on previous reports suggesting their association with fetal behavioral states ^42,43^. To uncover potential effects specific to certain vigilance states of the mother, the best-performing model was additionally applied to the subset of data corresponding to maternal sleep and wake only.

#### Longitudinal sleep behavior prediction

To analyze the infants’ sleep behavior across assessments, we sought to identify a common variable that could be extracted from all employed methodologies (i.e., abdominal acceleration signals during pregnancy, early postnatal diaries, as well as the Brief Infant Sleep Questionnaire at 3 and 6 months). Additionally, since the focus was on the maturation of sleep rhythms, this variable should be indicative of the child’s diurnal rhythms. Considering that the questionnaire provides information about the infant’s sleep duration between 7 pm and 7 am, as well as 7 am and 7 pm, we utilized these time windows to calculate the infants’ daytime and nighttime sleep duration across different time points. For this purpose, we considered all intervals within the defined time window labelled as fetal inactivity in abdominal acceleration signals and as sleep in early postnatal diaries. From the derived duration, a day/night sleep ratio was calculated for each infant and each time point. A random forest algorithm with 100 trees and 10-fold cross-validation was used to predict the early postnatal, 3-month, and 6-month day/night sleep ratio including the features outlined in Table 1. Procedures were repeated twice for each outcome, once including the gestational age at time of the outcome assessment and once including the chronological age at time of the outcome assessment. We then compared the models with regards to their mean squared error (MSE) to determine which measure of age is most relevant to the development of sleep rhythms. All continuous variables were standardized by means of z-scores within each fold on the training set only, to avoid data leakage from the test set. The resulting feature importance was averaged across folds to get a robust estimate. We sorted the features by their importance and calculated the cumulative sum to determine the number of features needed to exceed 95% of total importance. These features were interpreted as the best predictors. Finally, we retrained the model with only these features and visualized partial dependence of each feature to illustrate the association with the outcome variable.

**Table 1.** Input features modelled in a random forest algorithm with 100 trees and 10-fold cross-validation used to predict the early postnatal (first column), 3-month (second column), and 6-month (third column) day/night sleep ratio.

| **Feature** | **Early postnatal** | **3 months** | **6 months** |
| --- | --- | --- | --- |
| Fetal sleep ratio | x | x | x |
| Early postnatal sleep ratio |  | x | x |
| 3-month sleep ratio |  |  | x |
| Maternal temperature phase during pregnancy assessment | x | x | x |
| Maternal SRI during pregnancy assessment | x | x | x |
| Sex | x | x | x |
| Birth mode | x | x | x |
| Concurrent age | x | x | x |
| Concurrent breastfeeding (i.e., yes, no, partially) | x | x | x |
| Breast milk melatonin concentration | x | x | x |
| Infant stool melatonin concentration | x | x | x |

All analyses were conducted in MATLAB R2022b (Mathworks, Natick MA, USA) and R 4.2.2 ^44^ in RStudio 2022.12.0 (Posit PBC, Boston MA, USA).

## Results

At the start of the assessment, the participating women were between 20 and 40 years of age (mean age ± SD = 33 ± 4 years) and 195 to 271 days pregnant (235 ± 20 days). Twenty-six participants were nulliparous. Two pregnancies had resulted from *in vitro* fertilization, and one participant was diagnosed with gestational diabetes. Two of the pregnancies were twin pregnancies and, in these cases, the sensors were placed to ensure that movement from each twin could be distinctly monitored using two separate sensors. We only included data from the twin whose movements were more dominantly perceived by the mother. Further characteristics of the sample across time points are listed in Table 2.

**Table 2.** Distribution of the examined characteristics across the sample. Either the mean and the standard deviation with the range in parentheses or the number of observations with percentage in parentheses are shown for each characteristic.

| Characteristic |  |
| --- | --- |
| Fetal day/night sleep ratio | 1.11 ± 0.13 (0.81-1.33) |
| Gestational age at fetal assessment (days)<br>Mean ± SD (range) | 235 ± 20 (195-271) |
| Maternal sleep regularity index<br>Mean ± SD (range) | 0.67 ± 0.04 (0.61-0.8) |
| Maternal temperature phase (hours)<br>Mean ± SD (range) | 2.73 ± 1.01 (1.04-5.21) |
| Sex<br>N (%) | 18 females (58) |
| Early postnatal day/night sleep ratio | 0.73 ± 0.16 (0.5-1.04) |
| Age at early postnatal assessment (days)<br>Mean ± SD (range) | 22.1 ± 14.9 (2-54) |
| Breastfeeding at early postnatal assessment<br>N (%) |  |
| Yes | 10 (71) |
| Partially | 3 (21) |
| No | 1 (7) |
| Birth mode<br>N (%) |  |
| Vaginal | 13 (41) |
| Forceps | 2 (6) |
| Induced | 3 (9) |
| Caesarean | 12 (38) |
| Melatonin concentration in breast milk (pg/mL)<br>Mean ± SD (range) | 6.29 ± 6.59 (0-17.24) |
| Melatonin concentration in infant stool | 146.14 ± 101.02 (37.73-317.57) |
| 3-month day/night sleep ratio | 0.51 ± 0.18 (0.23-1) |
| Age at 3-month follow-up (days)<br>Mean ± SD (range) | 102.5 ± 9.9 (83-119) |
| Breastfeeding at 3-month follow-up<br>N (%) |  |
| Yes | 10 (75) |
| Partially | 4 (14) |
| No | 3 (11) |
| 6-month day/night sleep ratio | 0.27 ± 0.08 (0.16-0.44) |
| Age at 6-month follow-up (days) | 193.4 ± 9.8 (179-209) |
| Mean ± SD (range) | |
| Breastfeeding at 6-month follow-up |  |
| N (%) |  |
| Yes | 10 (24) |
| Partially | 15 (60) |
| No | 4 (16) |

Examples of processed abdominal signals along with the corresponding maternal perception of fetal movement from the diaries are shown in Figure 2. The Jaccard index, calculated as a measure of similarity between between maternal perceptions (i.e., reported fetal movement/missing report) and abdominal signal labels (i.e., fetal activity/inactivity), ranged from 0.25 to 0.7 with a mean of 0.42. This indicates a moderate level of agreement on average while revealing substantial individual variability. Notably, in 90% of cases, the sensor that showed the highest similarity to maternal perceptions corresponded to the maternally reported site of strongest fetal movement.

**Figure 2:**
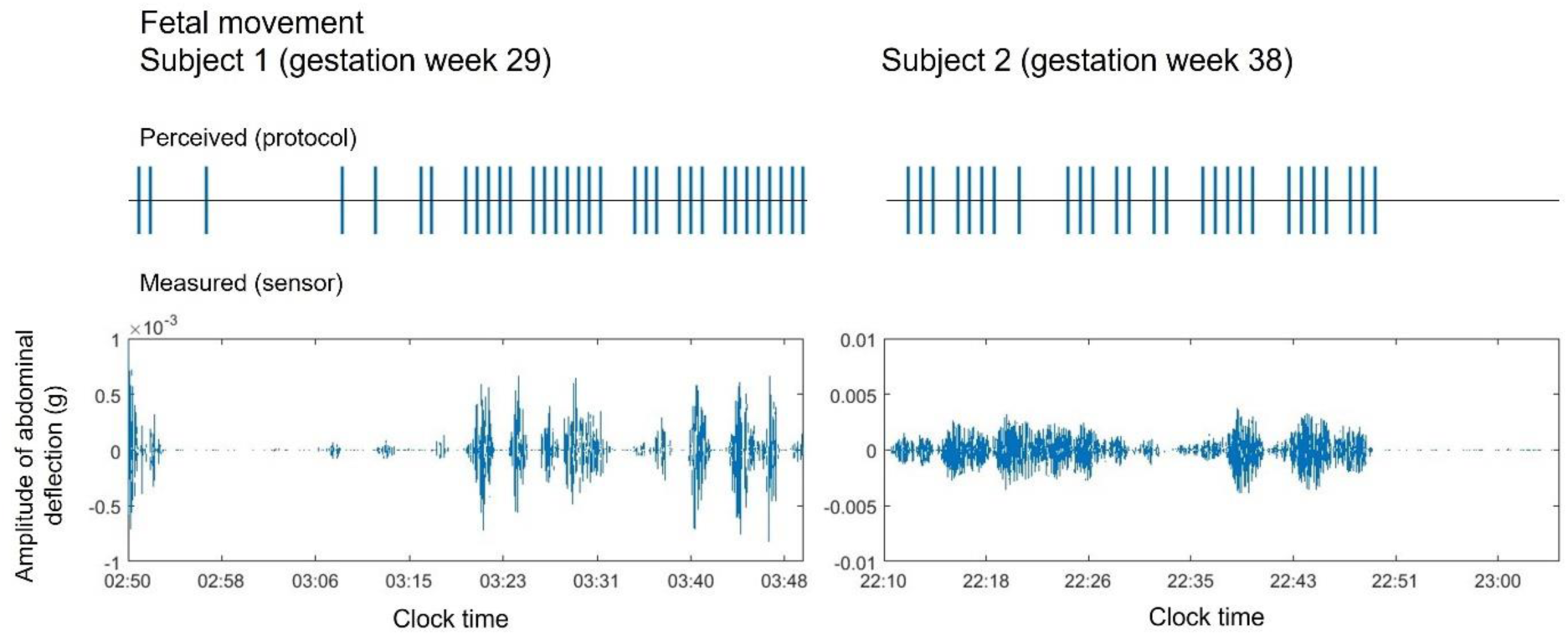
Two examples of fetal movement as perceived and documented in the minute-by-minute protocol by the mother during quiet rest (upper panels) and filtered acceleration signal amplitude from an abdominal sensor during the same time period (bottom panels). Lines in the upper panels depict protocol entries when the mother noted fetal movement. Amplitude of the signal in the lower panels reflects fetal movement, indicating that signals correspond well with the protocol.

### Pregnancy time-series modelling

#### Fetal activity is strongly associated with maternal rhythms

Binning the physiological signals from the pregnancy assessment to visualize hourly fluctuations within 24 hours as expected showed that maternal wrist skin temperature reaches its peak during nighttime, while wrist actigraphy-derived maternal activity is increased during daytime (Figure 3A). Fetal activity showed a similar pattern, with higher levels during daytime compared to nighttime (Figure 3A). Thus, both maternal and fetal activities displayed notable diurnal rhythms.

**Figure 3:**
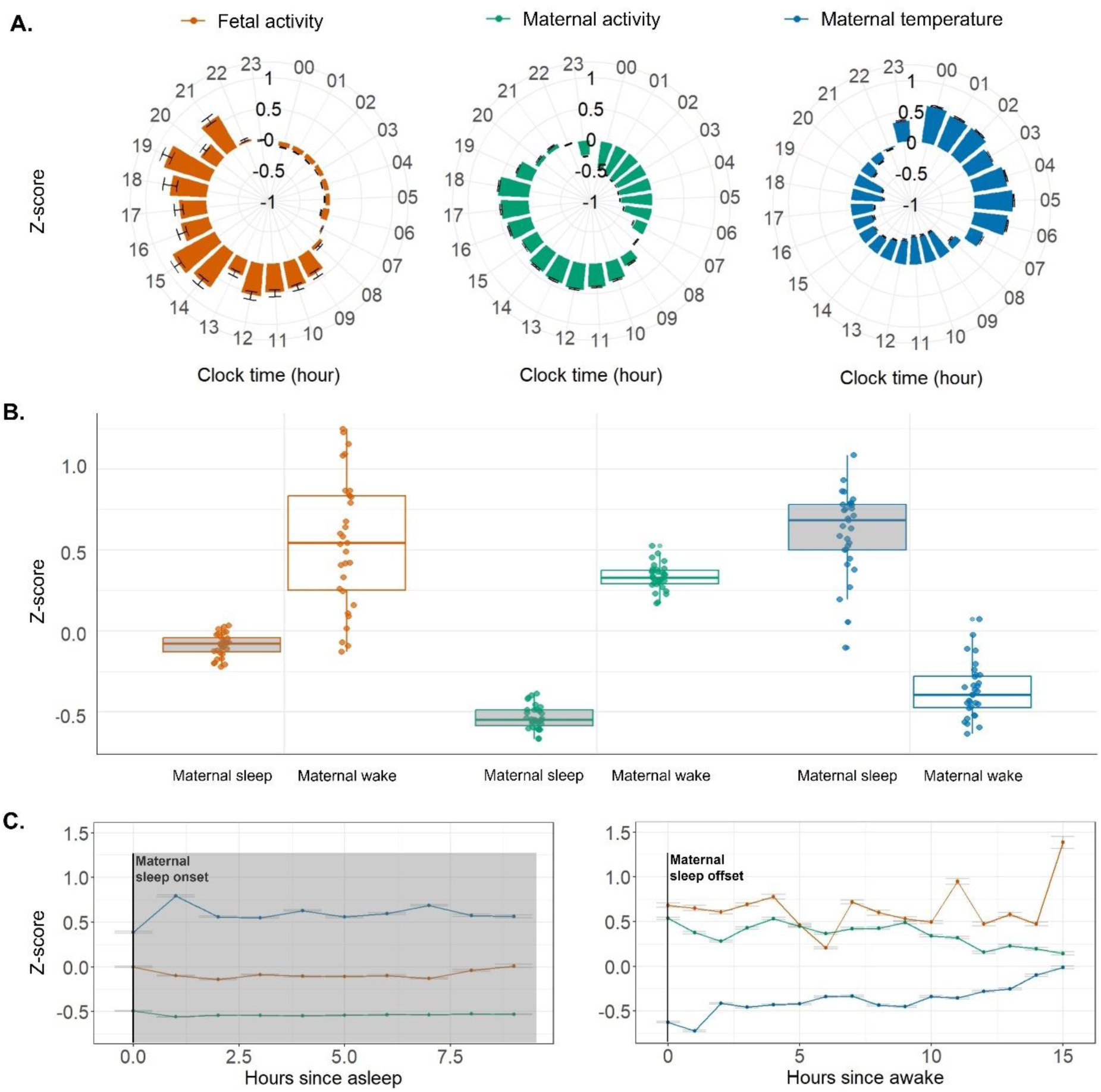
Z-scores of wearable-recorded signals in pregnancy, indicative of diurnal rhythms in fetal activity, maternal activity and maternal temperature. A) Diurnal profiles of signals averaged across 32 participants collapsed for the 24 hours of a day indicating a preference towards daytime activity. Larger numbers indicate increased activity or temperature as compared to the individual mean; B) Individual average signals in relation to the maternal vigilance state (i.e., periods of maternal sleep in gray shading and periods of maternal wake) showing a tendency towards fetal inactivity during maternal sleep, C) Mean trajectory of signals during maternal sleep (left panel with gray shading) and maternal wake (right panel) indicating diverging trajectories of fetal from maternal activity during maternal wake as opposed to sleep. The error bars depict the standard error.

The most complex multi-level model (i.e., model 4 including interaction terms) performed best (see Suppl. Table 1 for all models’ performance metrics). This model showed a main effect of gestational age, maternal vigilance state, maternal wrist skin temperature, and maternal wrist activity (Table 3). When stratified by maternal vigilance state, fetal activity during maternal sleep decreased with gestational age, whereas during maternal wake it increased with gestational age (Table 3). In other words, fetal preference towards nighttime inactivity increased with progressing gestational age (Figure 3B), possibly reflecting early development of diurnal rhythms. Similarly, fetuses were more active during periods of high maternal activity and low maternal temperature (i.e., during daytime; Table 3). However, when examining only periods of maternal wake, we found a negative association with higher levels of fetal activity being associated with lower levels of maternal activity (Figure 3C). Importantly, this observation suggests that fetal signals cannot be explained merely by maternal movement. The remaining factors’ effects were non-significant with the exception of nulliparity in the model including only data from periods of maternal wake (Table 3). In line with previous research ^45^, this association reflected lower fetal activity in nulliparous women. The remaining factors (i.e., twin pregnancy, in-vitro fertilization and gestational diabetes) did not show any significant effects.

**Table 3.**
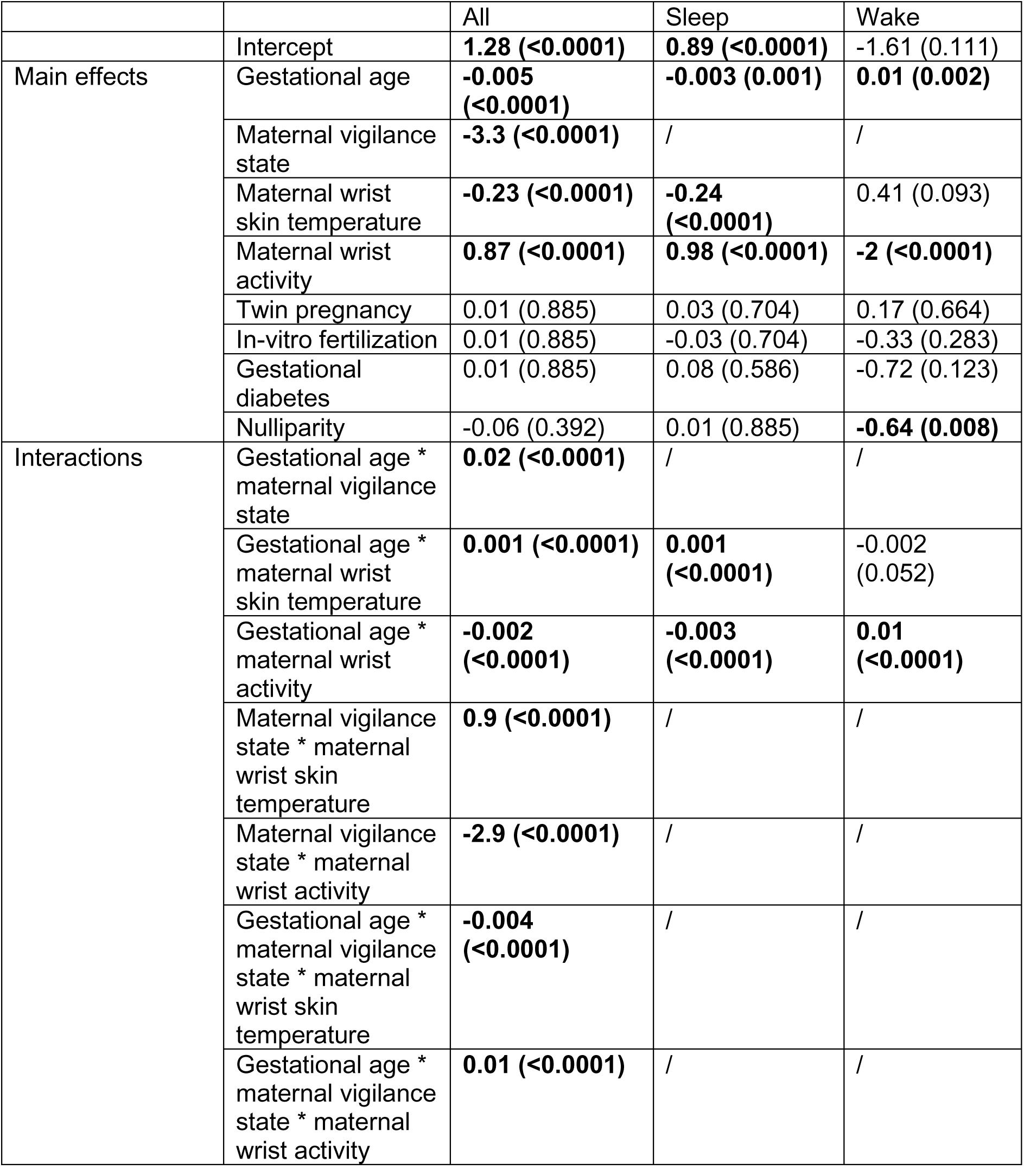
Results from multi-level regression model including physiological signals from 31 women during the last trimester of pregnancy. The main effects of gestational age (in days), maternal vigilance state (i.e., wake with maternal sleep used as reference), maternal temperature, twin pregnancy (yes/no), in-vitro fertilization (yes/no), gestational diabetes (yes/no) and nulliparity (yes/no), as well as interactions between gestational age, maternal vigilance state, maternal wrist skin temperature and maternal wrist activity with regards to changes in fetal activity signals (i.e., abdominal acceleration) are shown. The results from the same type of model are shown in the last two columns for the subset of data corresponding to maternal sleep or wake. In these models, the effects associated with maternal vigilance state are thus missing. Unstandardized beta coefficients are presented with p-values in parentheses corrected by means of the false discovery rate for the 36 conducted statistical tests (i.e., number of p-values across the three models). Significant effects are bolded (p < 0.05).

#### With age, fetal rhythms become increasingly independent of maternal rhythms

All investigated interactions among gestational age, maternal vigilance state, and maternal rhythms were significant (Table 3). Specifically, in further progressed pregnancies the association of fetal activity with maternal activity and maternal temperature was weaker (Supplementary Figure 1A). Specifically, for younger pregnancies (lowest gestational age tertile), an increase in maternal activity from two standard deviations was associated with an increase in fetal activity of 0.65 standard deviations. In contrast, for more advanced pregnancies (highest gestational age tertile), the same increase in maternal activity was associated with a smaller increase in fetal activity of 0.45 standard deviations. Similarly, an increase in maternal temperature by one standard deviation was associated with a larger decrease in fetal activity of 0.05 standard deviations in younger pregnancies as compared to a decrease in fetal activity of 0.01 standard deviations in more advanced pregnancies.

The association of fetal activity with both maternal activity and maternal temperature was stronger during maternal wake as compared to sleep, as demonstrated by the interaction with maternal vigilance state (Supplementary Figure 1B). During maternal sleep, an increase in maternal activity by four standard deviations was associated with an increase in fetal activity of 1.10 standard deviations, whereas during maternal wake, the same increase in maternal activity was associated with an increase in fetal activity of 3.30 standard deviations. Similarly, an increase in maternal temperature of two standard deviations was associated with a smaller decrease in fetal activity of 0.05 standard deviations during maternal sleep as compared to a decrease in fetal activity of 0.21 standard deviations during maternal wake.

These observations were confirmed in the subset only including periods of maternal sleep, where the association of fetal activity with both maternal activity and temperature became weaker with increasing gestational age. In contrast, the same association became stronger with gestational age when only data corresponding to maternal wake were included. Finally, the interaction between the maternal vigilance state and gestational age confirmed that older fetuses (i.e., the oldest tertile) were 2.3 times less active during maternal sleep and 1.7 times more active during maternal wake as compared to younger fetuses (i.e., the youngest tertile), further supporting the emergence of fetal rhythms and an early diurnal preference. Of note, when examining only the data corresponding to maternal sleep, the results remained similar to the full dataset suggesting that this subset of data is driving the overall results, likely due to a larger dataset.

### Longitudinal sleep behavior prediction

Next, we employed a random forest algorithm to test whether fetal and maternal rhythms during pregnancy predict sleep behavior during later development in infancy. We limit our presentation of results to models based on chronological age, as those including gestational age gave similar results while performing slightly worse.

#### Fetal rhythm is the strongest predictor of early postnatal sleep

The random forest algorithm yielded 5 features that together accounted for more than 95% of the total importance in the early postnatal day/night sleep ratio (Figure 4A; MSE=0.134±0.055): fetal day/night sleep ratio (31%), age at early postnatal assessment (27%), maternal SRI during pregnancy assessment (18%), sex (13%), and breast milk melatonin concentration (11%). Fetal day/night sleep ratio was identified as the most important predictor, indicating that fetuses with a stronger preference for nighttime inactivity tended to become infants with a stronger preference for nighttime sleep (Supplementary Figure 2A). Older age at the early postnatal assessment was associated with a lower concurrent sleep ratio suggesting a stronger preference towards nighttime sleep in older infants. Maternal SRI was the third ranked predictor with more regular maternal sleep behavior (i.e., less variability from day to day) being associated with a stronger preference for nighttime sleep in infants. Furthermore, female infants demonstrated a stronger preference for nighttime sleep as compared to males. Surprisingly, higher levels of average breast milk melatonin were associated with less pronounced preference for nighttime sleep in infants. Furthermore, we found a similar association when analyzing breast milk melatonin content using the intra-individual standard deviation of samples from the same mother as a proxy for circadian rhythmicity. In line with stool melatonin, we thus limit our report to average breast milk melatonin. In comparison to breast milk, infant stool contained remarkably higher concentrations of melatonin, with an average increase of over 20-fold (Table 3 and Figure 4D). The remaining variables (i.e., maternal temperature phase during pregnancy assessment, birth mode, breastfeeding at early postnatal assessment, and infant stool melatonin concentration) accounted for negligible levels of the total importance.

**Figure 4:**
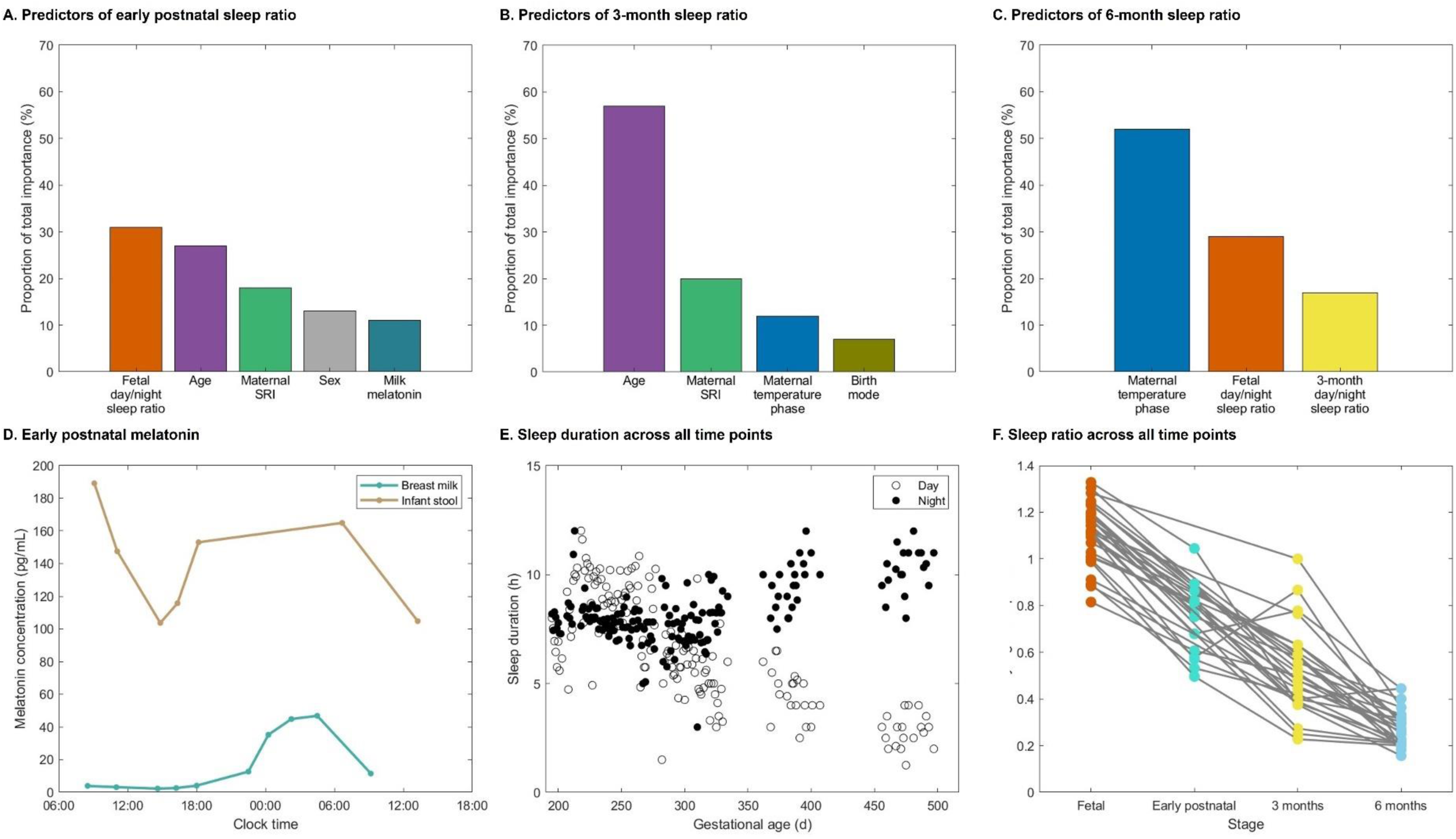
A-C) Results from random forest algorithm with 100 trees and 10-fold cross-validation. Set of predictors accounting for more than 95% of the total importance in the early postnatal day/night sleep ratio (A), 3-month day/night sleep ratio (B) and 6-month day/night sleep ratio (C). D) Exemplary plot of melatonin curves based on sequential data extracted from infant stool and maternal breast milk samples of one mother-child pair. E) Trajectory of sleep maturation showing all daily recorded sleep durations across time points as a function of gestational age. F) Trajectory of sleep ratio development, with data from the same individuals connected by lines. Lower ratios indicate a stronger preference towards nighttime sleep.

#### Regular maternal sleep in pregnancy predicts infant sleep at 3 months

Regarding the 3-month sleep outcome (i.e., day/night sleep ratio), 4 features together accounted for more than 95% of the total importance (Figure 4B; MSE=0.173±0.085): age at 3-month assessment (57%), maternal SRI during pregnancy assessment (20%), maternal temperature phase during pregnancy assessment (12%), and birth mode (7%). The exact age at the 3-month assessment was the most important predictor indicating a lower 3-month day/night sleep ratio in older infants, suggesting a continuing shift towards nighttime sleep (Supplementary Figure 2B). Similar to early postnatal models, a higher maternal SRI during pregnancy was associated with a stronger preference for nighttime sleep in 3-month-olds. Additionally, a later maternal temperature phase during pregnancy (i.e., maximal wrist skin temperature later after midnight indicating a later maternal chronotype) was associated with a lower 3-month day/night sleep ratio. Finally, birth mode accounted for 7% of the importance with infants born through a Caesarean section demonstrating a weaker preference towards nighttime sleep at 3 months.

#### Maternal rhythms in pregnancy and fetal sleep predict infant sleep at 6 months

Three features together accounted for more than 95% of the total importance in the 6-month day/night sleep ratio (Figure 4C; MSE=0.069±0.032): maternal temperature phase during pregnancy assessment (52%), fetal day/night sleep ratio (29%), 3-month day/night sleep ratio (17%). Similar to 3-month models, later maternal temperature phase during pregnancy was associated with a stronger preference for nighttime sleep at 6 months (Supplementary Figure 2C). Additionally, a stronger preference towards nighttime inactivity/sleep as demonstrated by a lower day/night sleep ratio at previous assessments predicted a stronger nighttime sleep preference at 6 months.

Daytime and nighttime sleep duration is shown for all datapoints across the study in Figure 4E, while the development of the sleep ratio across time points is depicted in Figure 4F. These visualizations illustrate a gradual shift from daytime to nighttime sleep in parallel with a decrease in total sleep duration with age. It is evident that this development starts before birth. Taken together, in addition to age and maternal rhythms, individual preference towards nighttime inactivity/sleep remains among the most important predictors of sleep behavior across assessments.

Consistent results were obtained when excluding data from the participant who used two abdominal sensors with a sampling rate of 12.5 Hz.

## Discussion

Using wearable technology, we investigated the earliest, prenatal, development of activity patterns through continuous monitoring of fetal activity. Our study shows that diurnal preferences are observable before birth and associated with maternal activity and circadian rhythms (i.e., wrist skin temperature). Furthermore, in a longitudinal within-subject design, we tracked the participants’ sleep behavior after birth and found that infant diurnal preference is strongly predicted by fetal diurnal preference, but also by maternal sleep regularity and maternal temperature phase during pregnancy. These insights reveal a potential for developing predictive models that could be instrumental in identifying infants at risk for sleep disturbances or developmental delays. Further, they pinpoint maternal sleep timing during pregnancy as a modifiable target for prenatal intervention supporting outcomes of infant sleep regulation postpartum.

### Fetal diurnal preference predicts infant sleep behavior

Uniquely, we monitored diurnal activity patterns of the fetus by means of continuous wearable-recorded signals across multiple days. By following the participants through their first weeks and months of life, we quantified predictors and gained insights into how prenatal behavior translates into infant sleep rhythms. In line with previous reports ^46^, we found a gradual shift towards nighttime sleep with increasing age. Crucially, we observed that the emergence of a human’s diurnal preference begins already during fetal development, aligning with studies in non-human primates that indicate oscillation of clock genes in the fetal suprachiasmatic nucleus, albeit with a smaller amplitude as compared to adult animals ^47^. Furthermore, we found that fetal rhythms robustly predict the postnatal rhythms, an observation that provides support for the practical relevance of the methodology we employed to monitor fetal activity. This postnatal continuity in diurnal preference agrees with previous research showing consistent patterns between fetal and infant behavioral states, although recorded over shorter time intervals (i.e., a few hours) ^11,12^.

### Maternal rhythms predict fetal and infant sleep behavior

Our physiological data confirm the association of maternal rhythms (i.e., sleep regularity and wrist skin temperature) during pregnancy with both fetal and infant sleep rhythms. Notably, one study in humans found that therapeutic suppression of maternal adrenal function inhibited fetal heart rate and limb movement rhythmicity during 2-hour cardiotocography recordings ^48^, highlighting the importance of maternal physiological signals as entrainment cues for fetal rhythms. Our data during the critical early postnatal developmental stage suggest that maternal sleep timing during pregnancy can affect their child’s postnatal diurnal preference. This extends a concept from animal studies to humans, as research in rodents indicates that chronodisruption during pregnancy, in the form of simulated shift work, leads to disrupted rhythmicity of circadian clock genes, hormones, and metabolites in the offspring ^49^. Our findings demonstrate the importance of maternal rhythmicity during pregnancy, with effects extending beyond the prenatal period. Incorporating maternal sleep regularity into prenatal care guidelines may serve as a preventive approach to foster healthy circadian rhythms in both mother and infant.

Postnatally, melatonin levels in breast milk were predictive of infant sleep behavior. Indeed, circadian rhythmicity has been shown to appear earlier in breastfed infants suggesting a role of breast milk melatonin as a facilitating factor ^50^. Surprisingly, we found lower levels of milk melatonin were associated with an increased preference for infant nighttime sleep. On the other hand, melatonin levels in infant stool samples were not associated with their diurnal preferences. This observation, so far uninvestigated in human infants, suggests that the digestive system may serve as a distinct source of endogenous melatonin. Based on findings from animal research, the gastrointestinal tract has been proposed as one of the leading extra-pineal sources of melatonin, with regulatory mechanisms and functional characteristics distinct from those of melatonin produced by the pineal gland ^51,52^. Indeed, our exploratory analyses revealed no relationship between breast milk and infant stool melatonin (Spearman’s rho = 0.12, p = 0.77; data not shown). Consequently, infant stool melatonin could be involved in pathways separate from those influenced by melatonin transferred through breast milk – an intriguing hypothesis that warrants further investigation.

### Postnatal environmental exposure supersedes maternal influence by 3 months of age

We identified the chronological age as more relevant than the gestational age with regards to infant diurnal preference across all assessments. This finding can be explained by the direct influence of the extrauterine environment after birth, providing additional external cues to support circadian development. Support for this notion is provided by previous research that demonstrated significant, yet partial, continuity in sleep patterns from fetal to early postnatal stage, suggesting opportunity for external influences ^11,12^. Crucially, in our study, early postnatal sleep rhythms were predicted by maternal sleep regularity during pregnancy. By 3 months, the infant’s sleep was predicted by both maternal sleep regularity and temperature phase during pregnancy. By 6 months, the infant’s sleep was primarily predicted by maternal temperature phase during pregnancy. As compared to sleep regularity, reflecting acute behavior, temperature phase is a reliable marker of the circadian rhythm, and thus largely genetically determined ^53^ and modulated by external signals such as light ^54^. Therefore, genetically influenced circadian components appear to play an increasingly significant role over time. Our observation may thus be explained by the infant’s chronotype that is masked during the early postnatal period yet becomes increasingly apparent from the age of 3 months. Moreover, environmental exposures after birth may supersede the influence of maternal behavior during pregnancy by the time the infant reaches 3 months of age, thereby reducing the impact of maternal sleep regularity before birth.

### Limitations

Even though maternal motion was accounted for, it is possible that our analysis of acceleration signals recorded from the abdomen does not fully disentangle fetal from maternal movement. Nevertheless, given the observed diverging patterns between maternal and fetal activity (e.g., during maternal wake), we are confident that our findings cannot be explained merely by maternal movement driving the fetal signals. Although the presented methodology to detect fetal activity through abdominal acceleration measurement requires further improvement, this approach harbours a great potential for future implementation in clinical obstetric diagnostics. As a next step, normative prenatal sleep rhythms need to be determined and compared to those observed in risk pregnancies. This will advance the technology for early detection of risks associated with fetal and infant development and rhythm disturbances, as early as in the womb. For future investigations of fetal sleep, we suggest considering the application of additional sensing modalities. These could include fetal heart activity monitoring ^55,56^, used in combination with actigraphy, to refine the distinction between fetal behavioral states. Finally, we acknowledge the modest sample size, particularly at the early postnatal assessment, that possibly left some of the effects undiscovered. However, considering the delicacy of this life stage, even small samples from this population yet with long-time monitoring (i.e. 20 000 hours of wearable technology data) hold significant value and offer unique insights into the earliest stages of development.

### Conclusion

The critical role of circadian function in health throughout the lifespan ^57^ and its influence on fetal development during pregnancy are increasingly recognized ^58^. Our study leverages a longitudinal design to trace developmental trajectories of sleep from pregnancy through early childhood, addressing the recent call for comprehensive developmental research on early predictors of circadian health ^23^. By innovatively employing wearable technology for fetal monitoring, we continuously characterize fetal diurnal activity patterns, advancing our understanding of prenatal circadian rhythms. Our findings demonstrate that the prenatal environment predicts postnatal sleep behavior and circadian regulation, and highlight actionable risk factors for preventive roadmaps. This points to a broader application of our findings in precision medicine, where understanding individual differences in circadian regulation could lead to tailored strategies for enhancing health outcomes in early life. For example, targeting vulnerable periods for fetal clock development could help mitigate future sleep issues and associated mental health challenges in both infants and parents. In light of modern challenges like artificial lighting and reduced daylight exposure, which increasingly disrupt circadian alignment, our findings underscore the potential benefits of early interventions, such as optimizing maternal sleep hygiene during critical periods, to improve postnatal sleep outcomes. Taken together, our study lays the groundwork for developing normative curves and risk profiles in clinics, as well as preventive healthcare recommendations, ultimately enhancing societal health and well-being on a broad scale.

## Acknowledgments

We thank the participating families for their valuable contributions to this study. We also acknowledge the teams at the involved institutions for their support and inspiring discussions.

## Author contributions

All authors reviewed and approved the final version of this manuscript.

Conceptualization: AM, SK; Data curation: AM; Formal analysis: AM; Funding acquisition: SK; Investigation: AM; Methodology: AM, SK, CM, CB; Project administration: SK; Resources: SK, PZ; Supervision: SK; Validation: AM, SK, CM; Visualization: AM, CM; Writing – original draft: AM, SK; Writing – review & editing: AM, SK, CM, CB, PZ.

