## Supplementary material for "From womb to crib: How fetal activity patterns *in utero* reveal postnatal sleep behavior"

Supplementary Table 1

| Model 1) Intercept-only | AIC | 242800 |
| --- | --- | --- |
|  | RMSE | 0.998 |
| Model 2) Fixed factors | AIC | 236400 |
|  | RMSE | 0.96 |
| Model 3) Random factors | AIC | 235700 |
|  | RMSE | 0.955 |
| Model 4) Interactions | AIC | **233900** |
|  | RMSE | **0.945** |

Accuracy metrics (Akaike Information Criterion [AIC] and root mean squared error [RMSE]) for each of the evaluated models. The model with the lowest AIC and RMSE was selected and used for further analysis (in bold).

Supplementary Figure 1

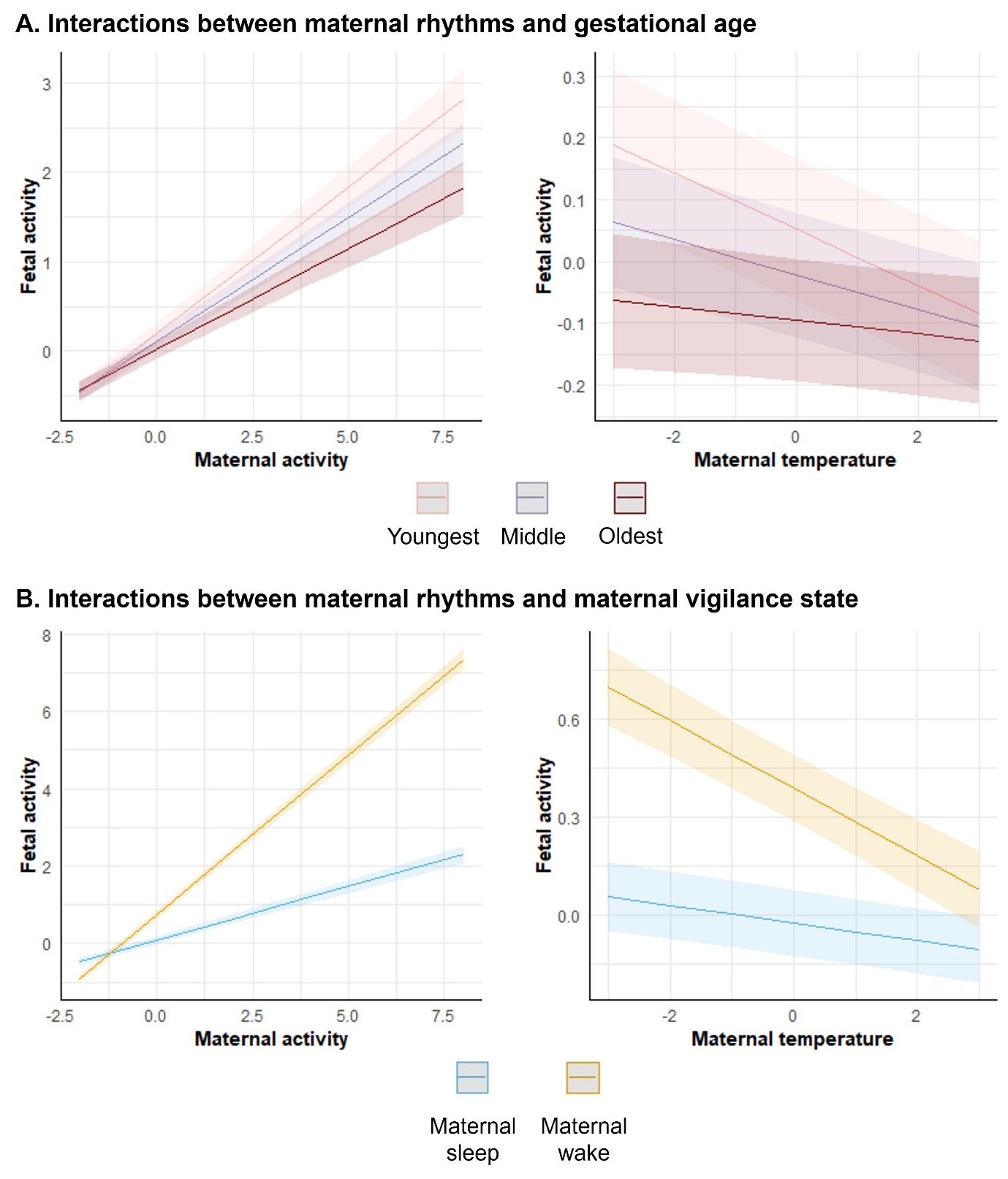

Interaction effects resulting from multi-level regression models including A) gestational age with the three age groups (i.e., youngest, middle and oldest) determined based on the tertiles of the age distribution within the study cohort, and B) maternal vigilance state (i.e., sleep and wake).

Supplementary Figure 2

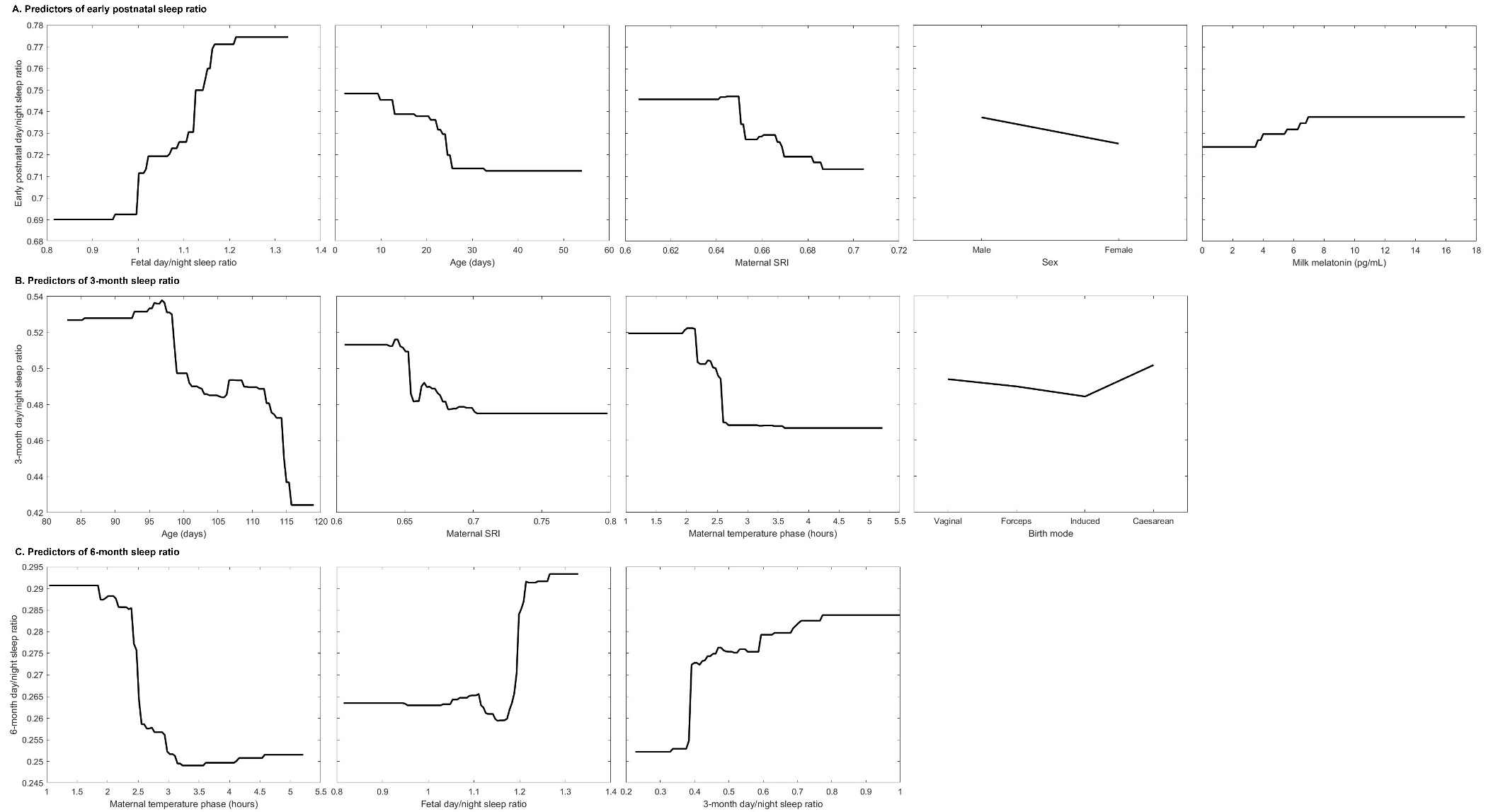

Partial dependence plots for each factor in the set of predictors accounting for more than 95% of the total importance in the early postnatal day/night sleep ratio (A), 3-month day/night sleep ratio (B) and 6-month day/night sleep ratio (C). Note that partial dependence plots are shown on the same scale for all predictors belonging to the same model (i.e., same outcome assessment). The y-axis in these plots represents the average predicted outcome of the model as that feature varies.
